# Modeling impairment of ionic regulation with extended Adaptive Exponential integrate-and-fire models

**DOI:** 10.1101/2024.08.01.606188

**Authors:** Damien Depannemaecker, Federico Tesler, Mathieu Desroches, Viktor Jirsa, Alain Destexhe

## Abstract

To model the dynamics of neuron membrane excitability many models can be considered, from the most biophysically detailed to the highest level of phenomenological description. Recent works at the single neuron level have shown the importance of taking into account the evolution of slow variables such as ionic concentration. A reduction of such a model to models of the integrate-and-fire family is interesting to then go to large network models. In this paper, we introduce a way to consider the impairment of ionic regulation by adding a third, slow, variable to the adaptive Exponential integrate-and-fire model (AdEx). We then implement and simulate a network including this model. We find that this network was able to generate normal and epileptic discharges. This model should be useful for the design of network simulations of normal and pathological states.

## 1 Introduction

In the study of seizure dynamics at the single neuron level, experiments have shown the importance of ionic regulation. Indeed, elevation of external ionic potassium concentration leads to seizure-like behaviors of the cells. Biophysical models have shown underlying mechanisms responsible for such activities [Depannemaecker et al., 2022b, Bazhenov et al., 2004].

Detailed biophysical network models have been developed to study seizure mechanisms [Rodrigues et al., 2015,Tejada et al., 2014,Santhakumar et al., 2005]. However, these models are complex with many parameters and variables, making their dynamics difficult to analyze. While seizures are typically seen as network events, similar dynamics can also be observed at the single-cell level in biophysical models [Bikson et al., 2003,Bragin et al., 1997, Chizhov et al., 2018, Cressman et al., 2009]. Phenomenological models, characterized by their minimal number of parameters and variables, enable comprehensive dynamical analyses and replicate many of the activities observed in experimental settings [Jirsa et al., 2014]. Some works integrate both approaches by reducing a detailed biophysical model to a lower-dimensional form, preserving the benefits of a generic model [Depannemaecker et al., 2022b].

However, these models are based on the Hodgkin-Huxley formalism [Hodgkin and Huxley, 1952], where the action potential occurs thanks to the nonlinear dynamics of the gating variables. The stiffness of these gating variables requires to use of very small time steps in order to simulate them accurately. This limits the use of such models as building blocks of large networks for long simulations. To address this issue we propose to reduce some aspects of the dynamics captured by the biophysical detailed model by using a formalism based on an integrate-and-fire type model. The adaptive exponential integrate-and-fire (AdEx) [Brette and Gerstner, 2005, Naud et al., 2008] model extends the classical integrate-and-fire (IF) model by incorporating two additional mechanisms: spike threshold adaptation through an exponential term and adaptation with an additional variable. In the AdEx model, the exponential term captures the subthreshold behavior of the neuron’s membrane potential. Specifically, the exponential term reflects the voltage-dependent conductance of the neuron’s membrane, which is a function of the difference between the neuron’s membrane potential and its threshold potential. The adaptation mechanism is modeled by an additional current term that reflects the neuron’s ability to adjust its firing threshold in response to the recent history of input. Specifically, this current term represents the activation of voltage-dependent potassium channels that contribute to the membrane’s afterhyperpolarization following a spike. The strength and time course of the adaptation current can be adjusted by the model’s parameters in order to reproduce a variety of observed neuronal responses. This adaptation variable enters as a current in the membrane voltage equation, which can create unrealistic values of individual membrane potential [Górski et al., 2021], but may not affect qualitatively the global dynamics at network levels [Depannemaecker et al., 2022a]. We thus introduce a third variable, which accounts for the effect of ionic impairment by modulating parameters that are conceptually associated with, or may be affected by ionic changes (e.g. reversal potential, spiking threshold).

Such a model can then become the basis to build large networks [Depannemaecker et al., 2022a], and hence study not only seizure propagation but also the emergence of such pathological patterns at the network level.

The article is organized as follows. In Section 2, we present both the single-neuron and the network model that we are going to analyze. Then, in Section 3, we show a correspondence with another model based on the Hodgkin-Huxley formalism and study different possible network configurations leading to the emergence of different patterns associated with epilepsy. We show how the severity of the local impairment or the size of the cell population concerned, leads to different patterns and propagation profiles. Finally, in Section 4, we summarise our findings and propose a few perspectives for future work.

## 2 Methods

We present here the single-neuron model. Subsequently, we explain how such a model is used for large network simulations.

### 2.1 Single-neuron models

The model builds on the AdEx [Brette and Gerstner, 2005, Naud et al., 2008], whose equations read:

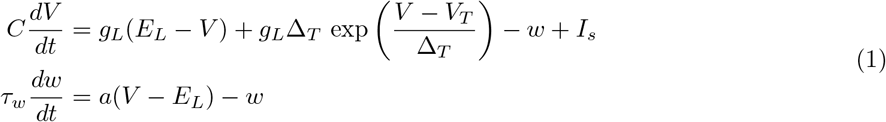

together with the following after-spike reset mechanism:

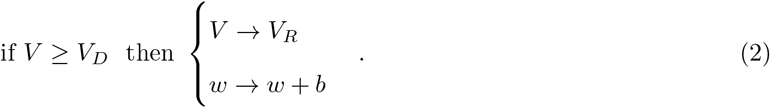

Based on this formalism, we added a third variable *z* with a slow timescale (with associated time constant *ε*) compared to the two other variables *V* and *w*. This variable *z* will phenomenologically aggregate the different impairments that may affect the ionic concentration regulation and thus the excitability of the neuronal membrane. Thus, this variable *z* modulates parameters (*E*_*L*_, *V*_*T*_), which enter into the other variables’ equations. The *E*_*L*_ parameter relates to the hyperpolarizing mechanism while the parameter *V*_*T*_ relates to the opening of sodium channels [Brette and Gerstner, 2005]. Hence, *V*_*T*_ may not be affected in the same way as *E*_*L*_, and it is thus scaled by parameter *β*. Furthermore, a counteractive term −*g*_*p*_*z* models phenomenologically the effects of the aggregate of biophysical mechanisms that are “working against the impairment” (e.g., the Na/K-pump, co-transporters, exchangers, etc…). Parameter *g*_*p*_ is the global equivalent conductance resulting from all these mechanisms. Finally, parameter *Z*_0_ corresponds to the potential to which the system will be attracted (depolarized) due to the pathological impairment. Therefore, the extended AdEx model that will study takes the form:

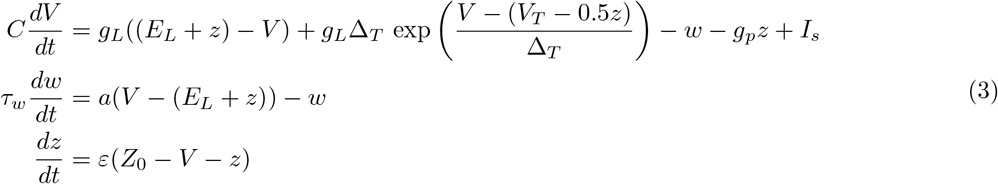

with still the after-spike reset mechanism:

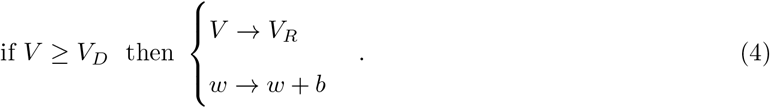

### 2.2 Network model

The network we will be studying is made up of inhibitory fast-spiking cells (FS) and excitatory regular spiking cells (RS). It contains 10,000 cells, 80% of which are excitatory and 20% inhibitory, connected through a random and sparse (Erdős-Rényi type) architecture with a probability of 5%. A conductance-based model of synapses enables connections between cells, it typically takes the form of equation (5) below:

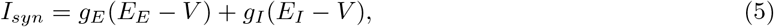

where *E*_*E*_ = 0 mV and *E*_*I*_ = −80 mV are the reversal potential of excitatory synapses and of inhibitory synapses, respectively. Parameters *g*_*E*_ and *g*_*I*_ are the excitatory and inhibitory conductances, respectively. They are governed by equation (6), and they are increased by a quantity *Q*_*E*_ = 1.5 nS and *Q*_*I*_ = 5 nS for each excitatory and inhibitory incoming spike, respectively. The timescale of synaptic conductance is determined by *τ*_*syn*_ = 5 ms.

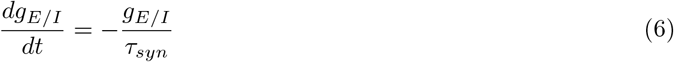

**Table 1:** Parameters values used (unless specified otherwise)

| Parameter | Value | Parameter | Value |
| --- | --- | --- | --- |
| $C_m$ | 200 pF | $g_L$ | 10 nS |
| $E_e$ | 0 mV | $a$ | 1 nS |
| $t_{\text{refractory}}$ | 5 ms | $b$ | 60 pA |
| $E_L$ | -65 mV | $V_T$ | -55 mV |
| $E_i$ | -80 mV | $I_s$ | 0 pA |
| $V_{\text{reset}}$ | -65 mV | $\tau_w$ | 0.5 s |
| $\varepsilon$ | $5 \cdot 10^{-4} \text{ s}^{-1}$ | $\Delta_T$ | 2 ms |
| $Z_0$ | -40 mV | $\tau_{\text{syn}}$ | 5 ms |
| $g_p$ | 10 nS | $V_D$ | -40 mV |

## 3 Results

Next, we present our findings in two distinct parts. First, we explore the behavior of our single-neuron model (3) in detail. This part will provide insights into how individual neurons function and react under various conditions, with a particular focus on the role of ionic regulation. In the second part, we delve into the dynamics of a large-scale neural network composed of 10,000 individual neurons. This investigation will help us understand how the characteristics of the extended AdEx model (3), discussed in the first part, influence the collective behavior and patterns that emerge within the network. This second part gives some insights into complex pathological phenomena such as seizure propagation.

### 3.1 Single neuron model

The single-neuron model exhibits different patterns of spontaneous pathological activities upon an increase of the value of *Z*_0_, consistent with what is observed in biophysical descriptions [Depannemaecker et al., 2022b]. However, the integrate-and-fire type model proposed here captures the after-spike repolarization via a reset mechanism. Thus, this after-threshold reset does not allow for the existence of a fixed point for values corresponding to depolarized membrane potential. Therefore, it is impossible to capture events with depolarization blocks. In this model, spontaneous activity appears for values of *Z*_0_ *>* 48 mV, and an increase of frequency is observed at the onset of a sustained tonic spiking pattern. While increasing *Z*_0_ to values higher than 40 mV, the model exhibits a pathological bursting pattern that can be associated with seizures. Finally, for *Z*_0_ *>* 20 mV the model dynamics corresponds to sustained ictal activity associated with status epilepticus and consisting of a constant high-frequency spiking pattern. After intense spiking activity, the current *w* reaches very high values leading to a strong negative input in the variable *V* that may take non-realistic values. This problem has been solved previously [Górski et al., 2021]; however, it may not affect the global network dynamics, specifically in the case of seizures as shown in a previous work [Depannemaecker et al., 2022a].

In order to understand the different dynamics observed in system (3), we considered *z* as being the slowest variable, enabling the loss of stability that leads to spiking activity. We then consider *z* as a parameter— which amounts to taking the limit *ε* = 0 in system (3) and obtaining its so-called *fast subsystem*— and we investigate the bifurcation structure of the resulting 2D system (with reset). The Jacobian matrix (7) of the fast subsystem is given by:

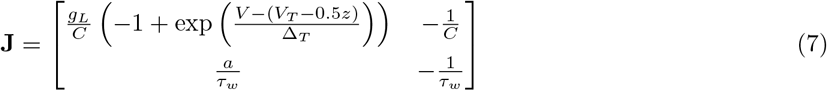

Hence, the trace of **J** takes the form:

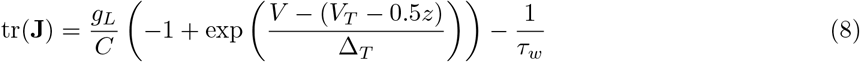

and its determinant reads:

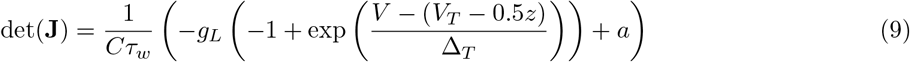

Thus, based on the trace and determinant, we can study the loss of stability in the *V* − *w* subsystem considering *z* as a parameter. In the figure 2 (a), we show the evolution of the two fixed points of the sub-system *V* −*w* (similar to the classical AdEx [Naud et al., 2008,Brette and Gerstner, 2005]). With the evolution of *z*, the spiking activity appears through a loss of stability by crossing an Andronov-Hopf bifurcation. The two fixed points then rapidly collapse and disappear, leading to the repeated spiking pattern. In the case of the pathological bursting pattern, the oscillation emerges from the interaction between the three variables *V*, *w*, and *z* (see figure 2 (b)). We thus separate two mechanisms leading to healthy or pathological bursts. Indeed, we should mention that the AdEx model is capable of generating bursting activity, as shown in detail in previous work by Naud et al. [Naud et al., 2008]. The original AdEx model requires an external input to spike (or specific parametrization that would lead to permanent steady-state spiking activity). In the model presented here, the spiking patterns appear through the interaction with the slow variable *z*.

**Figure 1:**
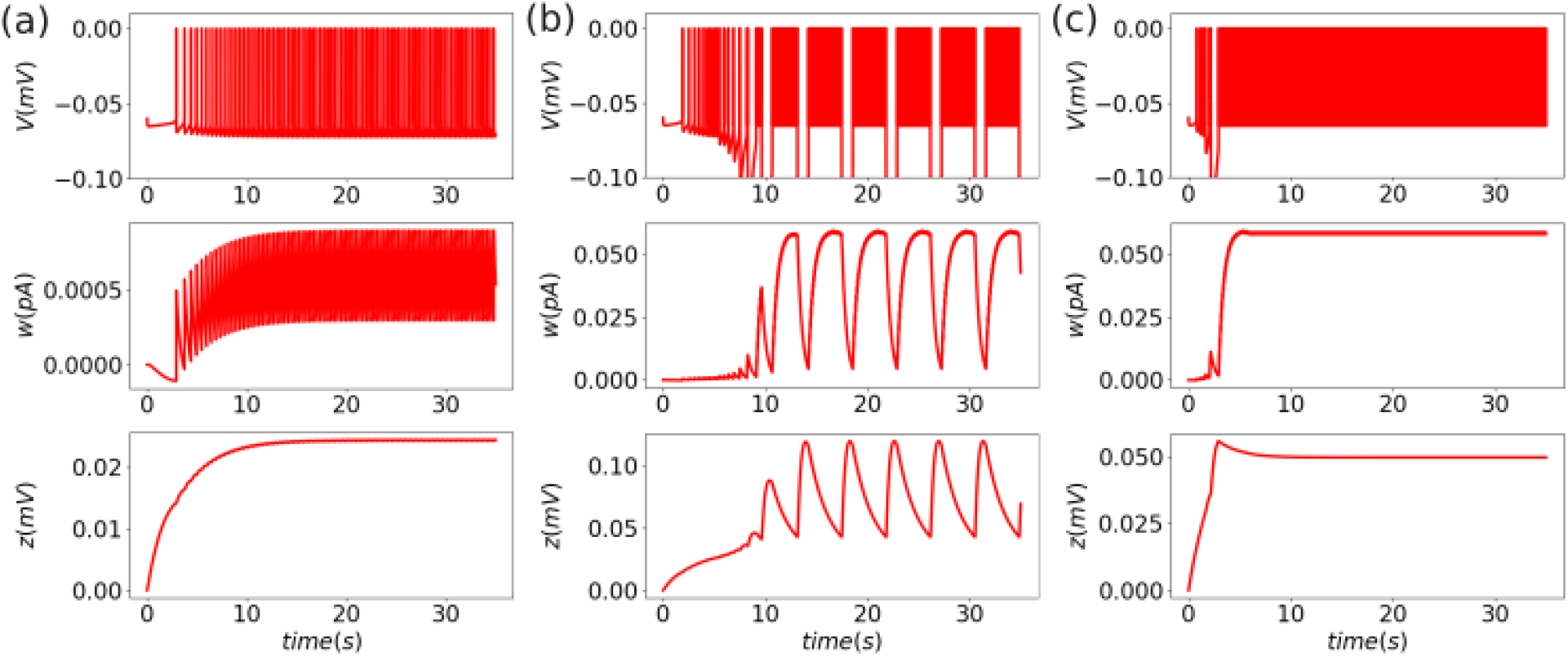
Spontaneous spiking patterns: (a) tonic spiking for *Z*_0_ = −45 mV, (b) pathological bursting pattern for *Z*_0_ = −40 mV (c) Sustain ictal activity for *Z*_0_ = −20 mV. The three simulations start from the same initial conditions, from which the (*V, w*) subsystem evolves to the specific pattern, driven by the effect of the slow variable *z*.

**Figure 2:**
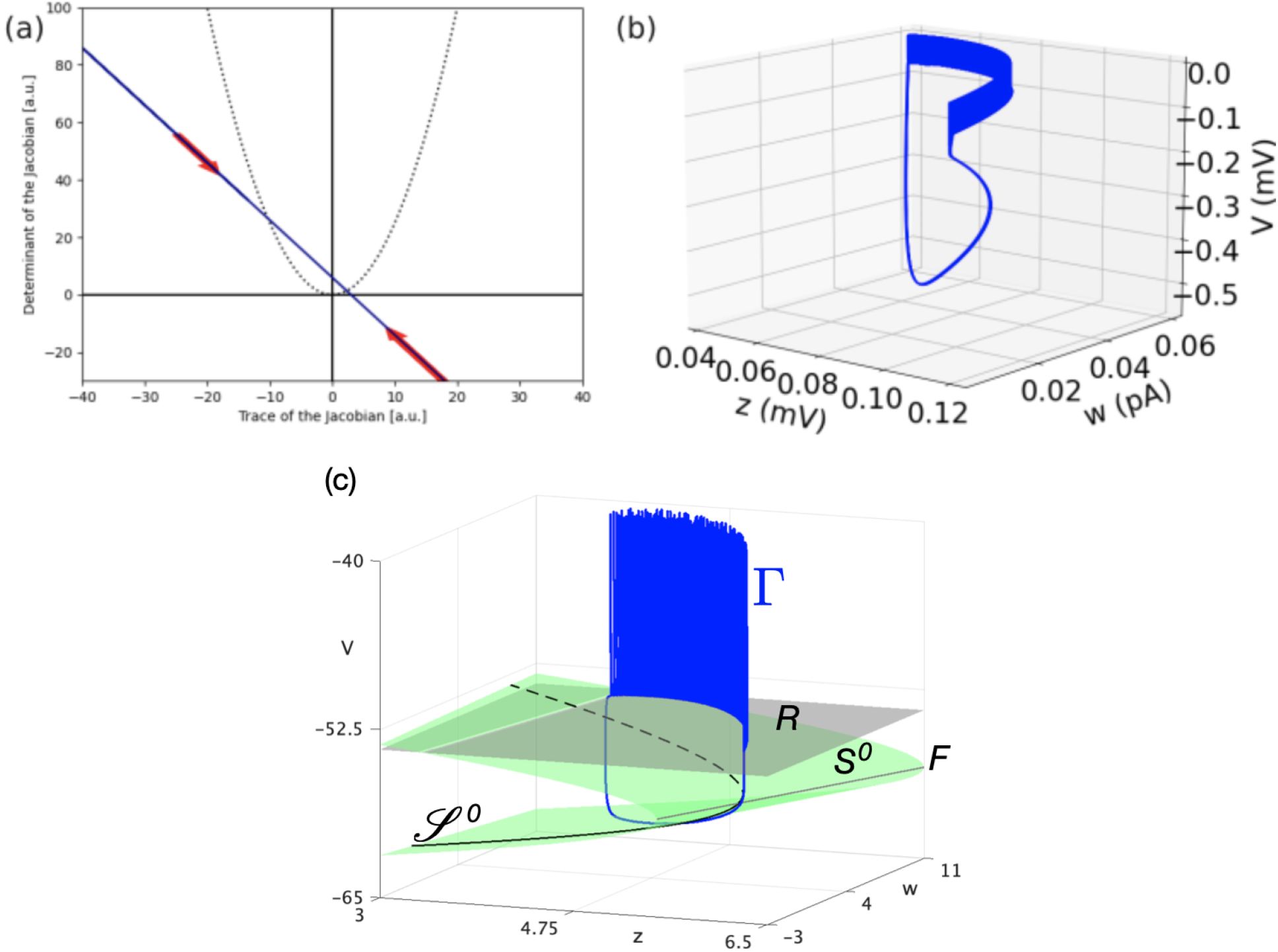
Bifurcations and trajectories: (a) Poincaré diagram of the fixed point of the *V* − *w* subsystem, the evolution of the slow variable *z* is considered as a parameter, the system loose stability crossing an Andronov-Hopf bifurcation (b) Pathological bursting pattern simulation plotted in the variable spaces, the trajectory does not exist in a single plane, the interaction between the three variables leads to this pattern. (c) Phase-space representation of another bursting pattern Γ (in blue), together with the Nullsurface of *v* (*S*^0^, in green, folded along the grey curve F), its intersection with the Nullsurface of *w* (𝒥^0^, in black), and the reset plane R. The associated parameter values are: *Z*_0_ = −51.2, *C* = 100, *g*_*L*_ = 10, *E*_*L*_ = −65, *α* = 1, *β* = 0.5, *a* = 1, *b* = 0.06, *V*_*D*_ = −40, *V*_*R*_ = −54, *I*_*s*_ = 0, *V*_*T*_ = −55, Δ_*T*_ = 2, *τ*_*W*_ = 200, *g*_*p*_ = 1, and *ε* = 0.001.

To understand the role of the slow variables we can plot the NullSurface into the phase space, and, project one trajectory of a simulation where the system exhibits bursting activity (see Fig. 2 (c)). We thus visualise the structure enabling the emergence of the bursting pattern.

The loop for bursting emerges with a concomitant effect of *w* and *z*, as shown by the trajectory in Fig. 2 (c), that exists in a surface in the diagonal on the *w* − *z* plan. The loss of stability at the onset of the burst occurs through a saddle-node bifurcation at the folding of the *v*-nullsurface (*S*^0^, in green in Fig. 2 (c)). The offset of the burst occurs through a (nonsmooth) homoclinic bursting, where the reset surface (grey) and the *v* Nullsurface (green) meet. The intersection of the *v*- and the *w*-nullsurfaces forms a curve 𝒥_0_ traced on *S*^0^, which the bursting dynamics would follow if *z* was the only slow process in the model. However, Fig. 2 (c) shows that the bursting pattern rather follows the surface *S*^0^ and much less the curve𝒥^0^. This indicates that *w* should also be considered as a slow process. Further analysis would be required to follow up on such considerations, which are an interesting topic for future work. For more elements about slow-fast bursting dynamics in IF models, we refer the reader to [Desroches et al., 2021, Desroches et al., 2024].

### 3.2 Network model

In this section, we detail how the severity of local impairment or the size of the affected cell population influences the emergence of distinctive patterns and propagation profiles in the context of epilepsy. To implement this analysis we performed a large parameter exploration, with a focus on the parameter *Z*_0_, which determines the level of local impairment, and the parameter *N*_*SC*_ which defines the number of impaired cells within the system. The results of the analysis are presented in Fig. 3 I. In panels a) to c) we show the maximum firing rates obtained during an 8-second simulation of the network as a function of *Z*_0_ and *N*_*SC*_. In panels d) to f) we show examples of the different dynamics obtained from the network. We see from this figure that the system’s behavior can be generally divided into three regions. In the first region (high *Z*_0_ and low *N*_*SC*_) the activity of the system remains in an asynchronous irregular (AI) regime with the impaired cells firing at a higher rate compared to healthy excitatory neurons but remaining within physiologically functional values. In the second region (intermediate values of *Z*_0_ and *N*_*SC*_), the firing rates of impaired cells exhibit already pathological dynamics with an oscillatory activity of large amplitude (up to 150Hz). In this second regime, the activity of the inhibitory cells follows the abnormal activity of the impaired cells, with the emergence of high-amplitude oscillations. This response of the inhibitory cells is enough to prevent the propagation of the pathological activity towards the healthy excitatory cells, which remain within normal values of firing rates with only moderate alteration in their dynamics. Finally, in the third region (low *Z*_0_ and high *N*_*SC*_), the pathological activity also propagates to the healthy excitatory population. For the example shown in panel f) of the figure we see that in the third regime, the entire system exhibits an alternation of periods of normal dynamics and periods of global increases in the firing rates. We notice that the pattern of activity observed in the third regime depends on the specific values of *Z*_0_ and *N*_*SC*_ and that the network model exhibits a large repertoire of possible patterns and dynamics. For completeness, we show in Fig. 3 II three different patterns obtained from different combinations of parameters.

**Figure 3:**
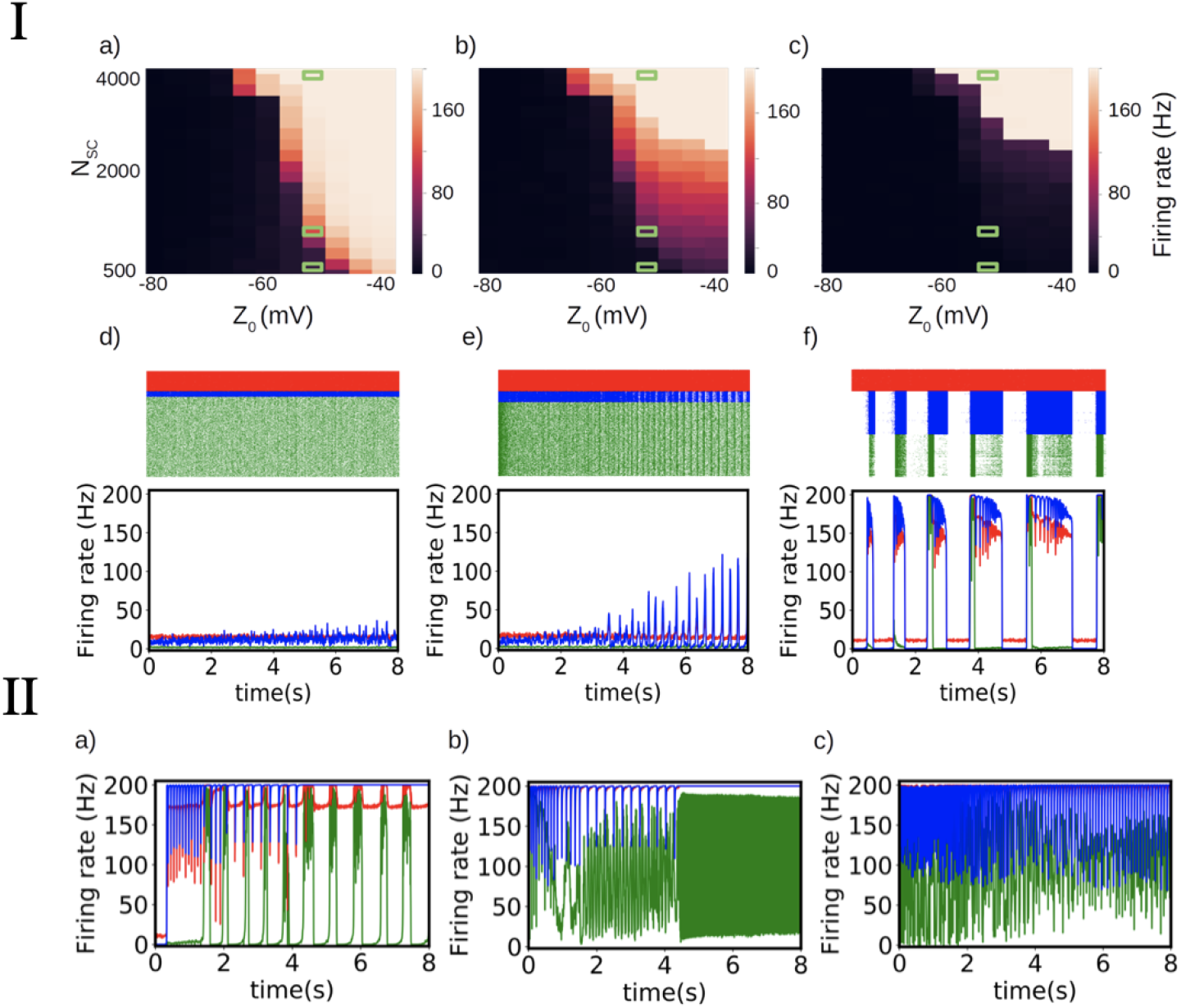
I-Propagation of pathological activity in a network model. a)-c) Maximum firing rates of the impaired cells (a), inhibitory cells (b), and excitatory cells (c) as a function of *Z*_0_ and *N*_*SC*_ (defining the severity of local impairment and number of impaired cells respectively. d)-f) Examples of different neuronal dynamics illustrate the effect of an increasing number of impaired cells *N*_*SC*_ leading to qualitative changes in the propagation of the pathological activity. For each case, the corresponding raster plot and average firing rate of each cell population is shown (impaired cells in blue, inhibitory cells in red, and excitatory cells in green). The plots correspond to the parameters *Z*_0_ = −50*mV* and *N*_*SC*_ = 500, 100, and 4000 respectively (the corresponding parameters are indicated with green rectangles in panels a,b,c).**II-Examples of different activity patterns for the propagation of pathological activity**. The network model exhibits a large repertoire of patterns of activity dependent of specific system parameters. We show in the figure the average dynamics obtained for three different sets of parameters: *Z*_0_ = −10*mV*, *N*_*SC*_ = 4000 (a); *Z*_0_ = −10*mV*, *N*_*SC*_ = 7500 (b) and *Z*_0_ = −40*mV*, *N*_*SC*_ = 7500 (c). The color code is the same as in I.

These network dynamics can be related to experimental observation [Wenzel et al., 2019]. Indeed, the propagation of seizure-like activities from one population of impaired cells to the rest of the network, follows a dynamics which resembles previous experimental observation. In Figure 3 we observe a threshold effect, where the propagation to healthy population is comparable to the phenomenon observable shown in Fig. 6 of the work of Wenzel et al. [Wenzel et al., 2019].

## 4 Discussion

Finally, this article has explored the dynamics of single neurons with an additional variable to account for ionic regulation impairment leading to pathological behavior. Experiments have highlighted the critical role of ionic regulation at the single neuron level, particularly in inducing seizure-like behaviors through elevated external ionic potassium concentrations. Biophysical models, although informative, have their limitations due to the Hodgkin-Huxley formalism, which necessitates small time steps and restricts their applicability in large network simulations [Cressman et al., 2009,Depannemaecker et al., 2022b]. To address this challenge, we have proposed an approach that simplifies certain aspects of the biophysical model using the adaptive exponential integrate-and-fire (AdEx) formalism, which includes subthreshold behavior modeling and adaptation mechanisms. This approach enables to capture different patterns emerging spontaneously from ionic changes. We show how the model captures these dynamics and their impact on neuronal responses. Furthermore, our exploration extends to the study of network configurations and their associations with epilepsy patterns, demonstrating how local impairment severity and cell population size influence patterns and propagation profiles.

If it is a simple model for seizure generation, it has some limitations in terms of dynamical repertoire. Indeed, a pattern associated with sustained depolarization (depolarization block, or permanent depolarization) can not be captured by this model due to the reset mechanism and thus the impossibility of the existence of a stable fixed point for high values (depolarized) of the membrane potential variable. In case of interest in these specific patterns, other models are more appropriate [Depannemaecker et al., 2023]. Such behaviors appear in mean-field approximation [Bandyopadhyay et al., 2021] of biophysically neuron models [Depannemaecker et al., 2022b].

Our approach proposed here is interesting to model large network dynamics and to capture the overall emergent dynamics. This simple model is thus a good candidate for building a mean-field model [Carlu et al., 2020, Alexandersen et al., 2024, Stenroos et al., 2024] capturing epileptic focus dynamics and which can be integrated into large-scale models of pathological brain states.

## 5 Acknowledgements

This research is supported by grants from the European Union, including the Human Brain Project H2020-94553 and the European Union’s Horizon Europe Programme under Specific Grant Agreement No. 101137289 (Virtual Brain Twin Project) and NSF-ANR (ImpactCom project). Additionally, funding has been received from the European Union’s Horizon Europe Programme under Specific Grant Agreement No. 101147319 (EBRAINS 2.0 Project). This work has also benefited from a government grant managed by the Agence Nationale de la Recherche (ANR) under the France 2030 program (PEPR), reference ANR-22-PESN-0012.

